# Image recognition based on deep learning in *Haemonchus contortus* motility assays

**DOI:** 10.1101/2021.12.01.470699

**Authors:** Martin Žofka, Linh Thuy Nguyen, Eva Mašátová, Petra Matoušková

## Abstract

Poor efficacy of some anthelmintics and rising concerns about the widespread drug resistance have highlighted the need for new drug discovery. The parasitic nematode *Haemonchus contortus* is an important model organism widely used for studies of drug resistance and drug screening with the current gold standard being the motility assay. We applied a deep learning approach Mask R-CNN for analysing motility videos containing varying rates of motile worms and compared it to other commonly used algorithms with different levels of complexity, namely the Wiggle Index and the Wide Field-of-View Nematode Tracking Platform. Mask R-CNN consistently outperformed the other algorithms in terms of the detection of worms as well as the precision of motility forecasts, having a mean absolute percentage error of 7.6% and a mean absolute error of 5.6% for the detection and motility forecasts, respectively. Using Mask R-CNN for motility assays confirmed the common problem with algorithms that use non-maximum suppression in detecting overlapping objects, which negatively impacts the overall precision. The use of intersect over union as a measure of the classification of motile / non-motile instances had an overall accuracy of 89%, indicating that it is a viable alternative to previously used methods based on movement characteristics, such as body bends. In comparison to the existing methods evaluated here, Mask R-CNN performed better and we anticipate that this method will broaden the number of possible approaches to video analysis of worm motility.

## 1. Introduction

Parasitic nematode infections present a substantial problem to human and veterinary medicine. Anthelmintics are often used as a weapon of choice to control or eliminate parasitic worms. With increasing reports of drug resistance, there is an ongoing effort to develop new anthelmintics [1, 2]. In view of this, drug screening and drug testing methods are clearly necessary. A number of important human and animal nematodes were already studied in large chemical library-scale screenings [3, 4], although nearly all of the studies were aimed at well-established non-parasitic *Caenorhabditis elegans* because of the scientific interest and medical applications as potential human disease models [5-9].

Nowadays, *in vitro* assays can measure the effects of compounds on development, growth, behaviour, and motility. Several approaches have been pursued in attempts to deliver robust, automated assays on the nematode phenotype [10-14]. The current gold standard for measuring drug effectiveness is *in vitro* assessment of worm motility, as measured via microscopy. The automated phenotyping from short video recordings offers a useful assay for high-throughput whole-organism phenotypic screening and avoids manual scoring. In assays with parasitic worms under experimental conditions, several challenges need to be overcome to enhance the performance of motility measurements. These challenges are associated with the different worm lengths, their tendency to clump, and also the range of diverse, complex movement patterns different from that seen in *C. elegans*.

Focusing on veterinary importance, the parasitic nematode *Haemonchus contortus*, known as the barber’s pole worm, is a model organism often used for drug screening and studying drug resistance; due to its economic importance, known genome and transcriptome [15, 16]. *H. contortus* infects predominantly small ruminants. It has a direct life cycle that starts with eggs being shed onto a pasture in the faeces of the infected animals. The cycle continues through three free-living larval stages to the parasitic fourth larval stage and lastly adults, which reside in the abomasum where they feed on blood. The ability to maintain *H. contortus* infection in a natural host animal and produce different larval stages *in vitro* provides experimental advantage. The infective third-stage larvae (L3s) can be stored, and researchers commonly use L3s or *in vitro* fourth larval stage for motility studies. Many automated or semi-automated approaches to measure motility have already been described such as via electrical impedance [17] or infrared light beam-interference [7]. Nevertheless, in our study of analysing the video recordings in *H. contortus* larvae to assess the motility, we have compared various (representative) algorithms with different levels of complexity in terms of model preparation as well as computational cost. We were interested to assess if the additional complexity of the methods leads to a sufficient increase in precision, that would justify the time-performance trade-off of these methods.

The first one mentioned is an algorithm which calculates a motility index value based on measuring the standard deviation of the pixel’s light intensity averaged for a number of frames [18]. This Wiggle Index (WI) does not allow for the detection of the phenotype of individual worms; still, this assay led to many “hit compounds” in screening of a large variety of compound libraries [19, 20]. The second approach called Wide Field-of-View Nematode Tracking Platform (WF-NTP) is built on the use of the Gaussian adaptive threshold of the image frames to identify worms and was applied in model free-living nematode *C. elegans* [21]. It was initially developed for drug screening purposes; nevertheless, it has also found applications in other studies such as detailed genetic or behavioural studies. Nonetheless, WF-NTP has a major limitation for images with overlapping objects as these objects get discarded by the algorithm based on an upper pixel size threshold per object, resulting in underestimation of the the total number of detected objects in an image [21]. In our study, we customized the associated WF-NTP software written in Python and compared it to the last approach which involves the application of deep learning.

The advent of machine learning has revolutionized the surveying and classifying of biological data including image recognition, enabling the automation of many tasks. The most important aspect of large-scale computerization is the possibility to automate *in vitro* assays and scale them, allowing researchers to avoid tedious repetitive tasks and focus on activities where they can use their knowledge to provide added value, leaving the analysis and interpretation of the data as the only rate-limiting step. In the field of computer vision, deep learning algorithms with a convolutional neural network (CNN) architecture have made rapid advancements on a variety of image classification tasks in cell cultures [22], and the model organism *C. elegans* [23, 24], even parasitic nematode in plants [25], yet none of them have been applied to parasitic gastro-intestinal nematodes of livestock, including *H. contortus*. Object-detection-based deep learning methods have also been adapted for instance segmentation. Here we used a state-of-the-art region-based CNN called Mask R-CNN [26], which can be trained based on a set of annotated images (Fig 1a) to detect the objects of interest, in our case the larval stages of *H. contortus*. In the first step, the algorithm uses a regional proposal network (RPN), i.e., a binary classifier that identifies rectangular subsets of the image that are of interest for the given task and are called regions of interest (ROI). In our case that would be parts of the image that potentially contain worms. These ROIs are then refined to cover the whole object (Fig 1b) and filtered using non-maximum suppression (NMS). NMS is a process of removing highly overlapping ROIs, to avoid detecting the same object multiple times before passing the filtered ROIs as proposals for classification. The classifier performs a refinement of the ROI and outputs a final rectangle surrounding the detected object called a bounding box, the class of the predicted instance along with the detection confidence and the object mask, which are the pixels that make up the object (Fig 1c).

**Fig 1:**
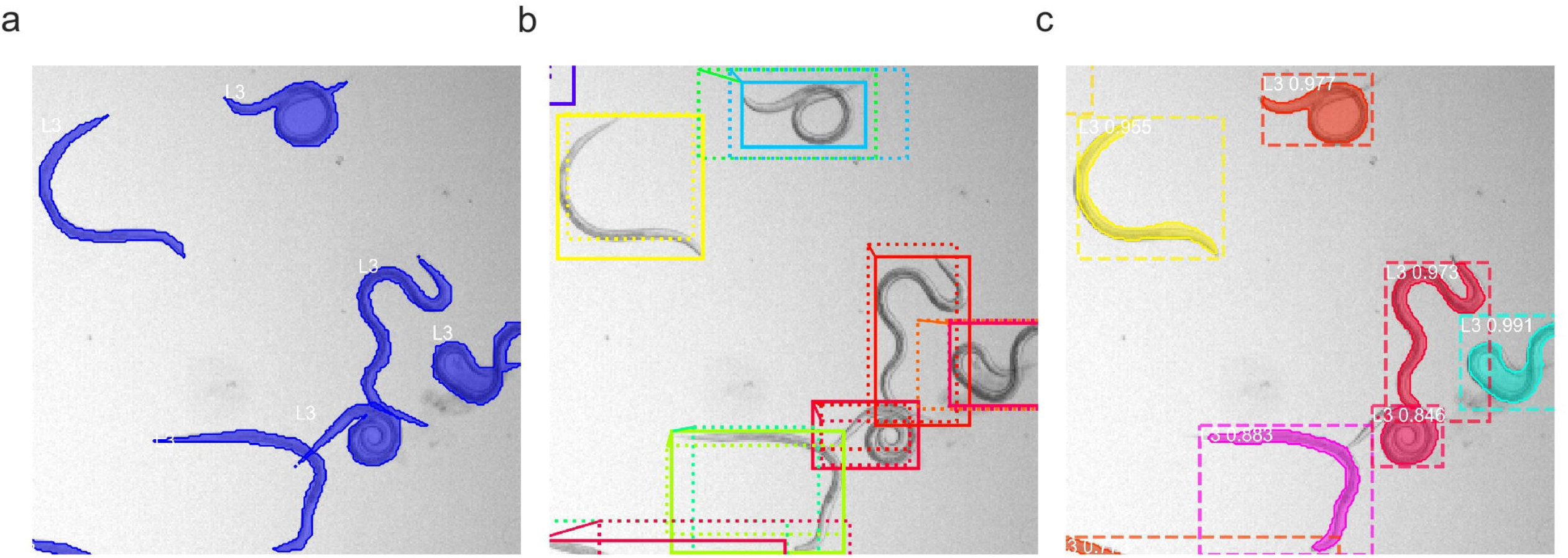
Mask R-CNN training and detection process **(a)** annotated images used for training the algorithm, **(b)** regions of interest (ROI) detected in the first step by the regional proposal network (solid boxes) and the refined regions (dashed boxes), **(c)** Mask R-CNN predictions containing the object boundaries, predicted object classes (i.e., L3 – third-stage larva) with an associated confidence level and the mask of the object (coloured contour).

The aim of the study was to train a Mask R-CNN model for image recognition of different life cycle stages of *H. contortus* and evaluate the application of this deep learning algorithm to the field of drug discovery of anthelmintics. Thereby, we compared the Mask R-CNN detections and motility predictions to the WI and WF-NTP methods.. Additionally for the motility, we tested a metric called intersection over union (IoU) instead of body bends from WF-NTP readout. IoU is used regularly in image recognition to quantify the extent of overlap between two objects, the intersection of the objects is divided by the union of the objects, which results in a numerical value between 0 and 1 (see Graphical abstract). Furthermore, we discussed the possible application of automated Mask R-CNN in other assays related to the veterinary practice.

## 2. Materials and methods

### 2.1. Procurement of L3s of *H. contortus*

Six-month-old sheep were orally infected with 8,000 infective L3s of MHco3 strain (Inbred Susceptible Edinburgh, ISE) [27]. Four weeks post-infection faecal samples were collected. L3s were produced from eggs by incubating faeces in a plastic box covered with foil at 27 °C for 7 days. Then, the faeces were rinsed twice in tap water which was poured into 1,000mL conical measuring cups, in which the larvae sank to bottom. To remove dirt or dead individuals, the pellet of larvae was pipetted out and filtered through a 20μm sieve submerged in water (27 °C). Clean L3s at a concentration of approximately 4,000 L3 per mL were stored in culture flasks in water at 10 °C for several months.

During the storage, some larvae naturally die. Prior to the experiment, we filtered larvae through a 20μm sieve (for 12 h), this time collecting both live larvae from the bottom of the sieve and dead individuals remaining on the sieve.

All experimental procedures were examined and approved by the Ethics Committee of the Ministry of Education, Youth and Sports (Protocol MSMT-25908/2014-9).

### 2.2. Video processing and manual counting

Videos from a microscope camera were recorded for a duration of 10 seconds with a framerate of 20 frames per second (fps) with a pixel quality of 2560 by 2160 (see Supplementary Video). The videos (.avi with MJPEG compression, ∼ 0.45GB) were acquired using NIS-Elements Imaging Software (version 4.20) using a Nikon Eclipse Ti microscope (4x magnification) with the camera (Andor Zyla 5.5 sCMOS, 12bit: 2560 × 2160 pixels). The videos were recorded for different ratios of live and dead larvae (100:0, 75:25, 50:50, 25:75, 0:100; denoted as motility group 100, 75, 50, 25 and 0, respectively) in 9 replicates and they contained on average 60 worms. The live larvae were kept at 37 °C to maintain their motility.

The video recordings were processed so that each worm was labelled with a numeric identifier. A trained human with prior experience of manual motility assays evaluation assessed each worm from each video and information about motility and the dead phenotype was recorded. The discrepancies were dealt with as follows (i) a dead worm which was moving due to the interference of a motile worm was counted as motile; (ii) a live coil non-motile worm was counted as non-motile; (iii) the worms which were not labelled by an algorithm were given an identifier manually; (iv) worms were excluded from the counts when they appeared in less than 20 frames; (v) dead phenotype was recorded when overt microscopical pathologies, such as the presence of large vacuoles in the tegument and straight phenotype, were observed.

This manual counting was taken as the referential method for comparison of accuracy among the three models.

### 2.3. Wiggle Index analysis

We used an ImageJ implementation of the WI for the analysis [28]. We did not apply any scaling of the videos, the Gaussian image blur was set to 2, the number of frames to be averaged was set to 50 and the motility output was based on the standard deviation. Since the values from the WI are relative, we used motility group 100 as the control group to be able to convert these relative values to percentages. We calculated the mean value for motility group 100 and divided all values by the mean to obtain normalized motility values.

### 2.4. WF-NTP analysis

WF-NTP presents an automated evaluation of motility in *C. elegans*. For its successful implementation, we adopted the method for the *H. contortus* motility assay. The algorithm was based on three steps;in the first instance all worms in each frame were detected, in the second step, these detections were linked based on a neighbourhood search, and in the final step, each worm was classified as motile or non-motile based on bends per second and the speed of the worm. However, as these parasites differ in their sizes as well as their movement, some modifications to the original configuration and algorithm were necessary. These modifications belonged predominantly to two categories, the first was a recalibration of the available input parameters of the WF-NTP method. This included aspects such as the image size, the size of the worms that we want to detect and other configurations (see Data availability). The other modifications were changes in the evaluation of the conditions for worm motility and a limit on the minimum number of frames that a worm needed to be detected in for it to be included in the evaluation.

The original method relied on bends per second and the speed of individual worms to classify them as either motile or non-motile [21]. When a worm was below a certain threshold of bends per second and below a certain speed threshold, it was classified as non-motile. In our settings, the speed threshold was obtained from the motility group 0. From the initial test for the motility group 50, the result was that out of 169 worms classified as motile, 23% exceeded the speed threshold, but no bends were recorded. This failure to measure bends per second for motile worms, was most likely caused by the more complex movement of *H. contortus*., This led us to replace the original approach with an evaluation metric, IoU, that is regularly used in image recognition [29]. The use of IoU allowed us to use the capabilities of WF-NTP in terms of the detection and tracking of individual worm instances in an image and then apply the IoU metric to determine the worm motility. The calculation of IoU was done for the same worm instance in neighbouring frames to detect changes on a frame-by-frame basis. Afterwards, the mean was calculated and worms above a certain IoU threshold were labelled as non-motile. To ensure that the comparison of WF-NTP and Mask R-CNN was aimed at the quality of the detection of individual worm instances rather than the approach to determine the motility, we used the same tracking for detections across frames and the same IoU conditions for both methods.

Another modification made to the original method was the introduction of a threshold for the minimum number of frames that a worm needed to be detected in, in order to be included in the analysis. The reason for this restriction was that as the individual worms move, they have a tendency to change shape or overlap with other worms, therefore in terms of the detection the individual worm can be lost for a certain number of frames and reappear later, however this can lead to counting some instances multiple times. To limit these potential duplicates (Supplementary Fig S1) as much as possible and not bias the total number of worms detected in the video, we selected the highest number of worms detected in a given video across all frames and added a buffer of 10% in order to account for situations where some worms were not detected, for example due to their overlaps. This maximum number then served as a limit for selecting the number of worms to be evaluated as motile or non-motile. The instances were selected based on the number of frames in a decreasing order, therefore ensuring that we had data from enough frames to classify the worm correctly. Again, this restriction was applied to detections from Mask R-CNN to enable a comparison under the same conditions.

### 2.5. Mask R-CNN analysis

For Mask R-CNN we used the Matterport implementation (https://github.com/matterport/Mask_RCNN) of the algorithm, including a trained model using a backbone architecture of Resnet101. The pre-trained model weights were obtained from training the model on the MS COCO dataset [30]. The process of using a pre-trained model is called Transfer Learning, which is an approach where the knowledge gained from solving one problem is applied to a related problem. The motivation for selecting a pre-trained model is that in general deep neural networks require a lot of annotated data for training and using an existing model and re-training the top layer of the network allows us to have a smaller dataset while maintaining good performance. We manually annotated a total of 95 images and divided them into training (69) and validation (26) using the VGG (Visual Geometry Group) Image Annotator [31], these images contained three categories of objects; eggs, first-stage larvae (L1), and L3, to fully utilise the capabilities of Mask R-CNN which is able to do instance segmentation and classification. The total number of annotated objects was around 10,400 divided into training (6,800 objects) and validation (3,600 objects). Then we ran the training on a GPU instance until the loss function showed no signs of improvement. To avoid overfitting, we used data augmentation and applied one or multiple of the following augmentations: flipping, contrast normalization, additive gaussian noise, multiplying the pixels by a number within a given interval to make the whole image lighter or darker. The final model had a mean average precision at 0.5 IoU of 64.1% on the validation set.

The trained model was then used on the individual frames of the video to detect the instances of parasites and we calculated the centroid of the detected object in terms of the position on the x and y axis using the scikit-image regionProps functionality [32]. These detected instances were then linked together using the trackpy library [33], which detected the trajectories of objects based on certain conditions. In our case we restricted the distance that the centroid could move a maximum of 100 pixels in each dimension. As a result of detecting the trajectories, we were able to identify the same parasite instance across multiple frames and assign it with a unique identifier. In cases where the tracking did not meet the above conditions, a new identifier was assigned, and therefore a single worm could be detected multiple times, which occurred for complex cases with extensive movements and a high number of overlaps. Subsequently, we calculated the IoU and the threshold for the number of parasites to select for the evaluation as described in the WF-NTP section. We observed a decreasing mean percentage error (MPE) with an increase in the length of the video, and therefore we decided on using the full length of the videos for the motility assay (Supplementary Fig S2).

### 2.6. Error metrics

For the evaluation of the performance of the individual algorithms, we used standard metrics for the quantification of the forecast precision. These metrics work with the error term, which is defined as the difference between the actual and the forecasted value.In our case, the actual value was obtained by manually processing the videos. Two aspects were measured; the first one was the bias of the forecasts to determine if the algorithm is systematically over or under-estimating, for that purpose, we used mean error (ME) or mean percentage error (MPE), which both work with the mean of the error term. The second aspect was the size of the error term irrespective of the error term being positive or negative to compare the overall performance of the algorithms. For that purpose, we used mean absolute error (MAE) or mean absolute percentage error (MAPE), which both work with the mean absolute value of the error term.

### 2.7. Monte Carlo drug screening simulation

To assess the impact of the differences in detection accuracy we ran a Monte Carlo drug screening simulation [34] for all three algorithms. To remove the effect of individual motility groups we worked with the differences between the predicted motility rates and the manually obtained motility rates. Using these differences we were able to obtain the correlations between the algorithms as well as the means and standard deviations. A multivariate normal distribution was used as the data generation process for the simulation to ensure that the correlation among the algorithms was taken into account. The SciPy package (version 1.3.3) implementation of the multivariate normal distribution was used.

For the simulation, we generated 1 million random motility rates from a continuous uniform distribution within the range of zero to one. Similarly, we generated 1 million observations of the multivariate normal distribution. These observations were the detection errors of each of the algorithms. By adding this error to the randomly generated motility rate, which served as our true observation, we received simulated motility predictions from the three algorithms. A “hit” in our simulation was defined as having a motility rate lower than 50%.

## 3. Results

### 3.1. Detection of motility

The observed typical cases for detection scenarios in the motility videos were depicted in Fig 2, where the results of Mask R-CNN were used for illustrating these examples. The most common situation, which did not pose a problem to the algorithms, were worms with few or no overlaps (Fig 2a). Other common occurrences were collisions between worms that caused a non-motile worm to be moved and therefore being perceived by the algorithms as motile (Fig 2b). Lastly, the most challenging scenarios contained a high number of worms with significant overlaps with mostly motile worms, which resulted in problematic detections and negatively impacted the ability to track worms across multiple frames (Fig 2c).

**Fig 2:**
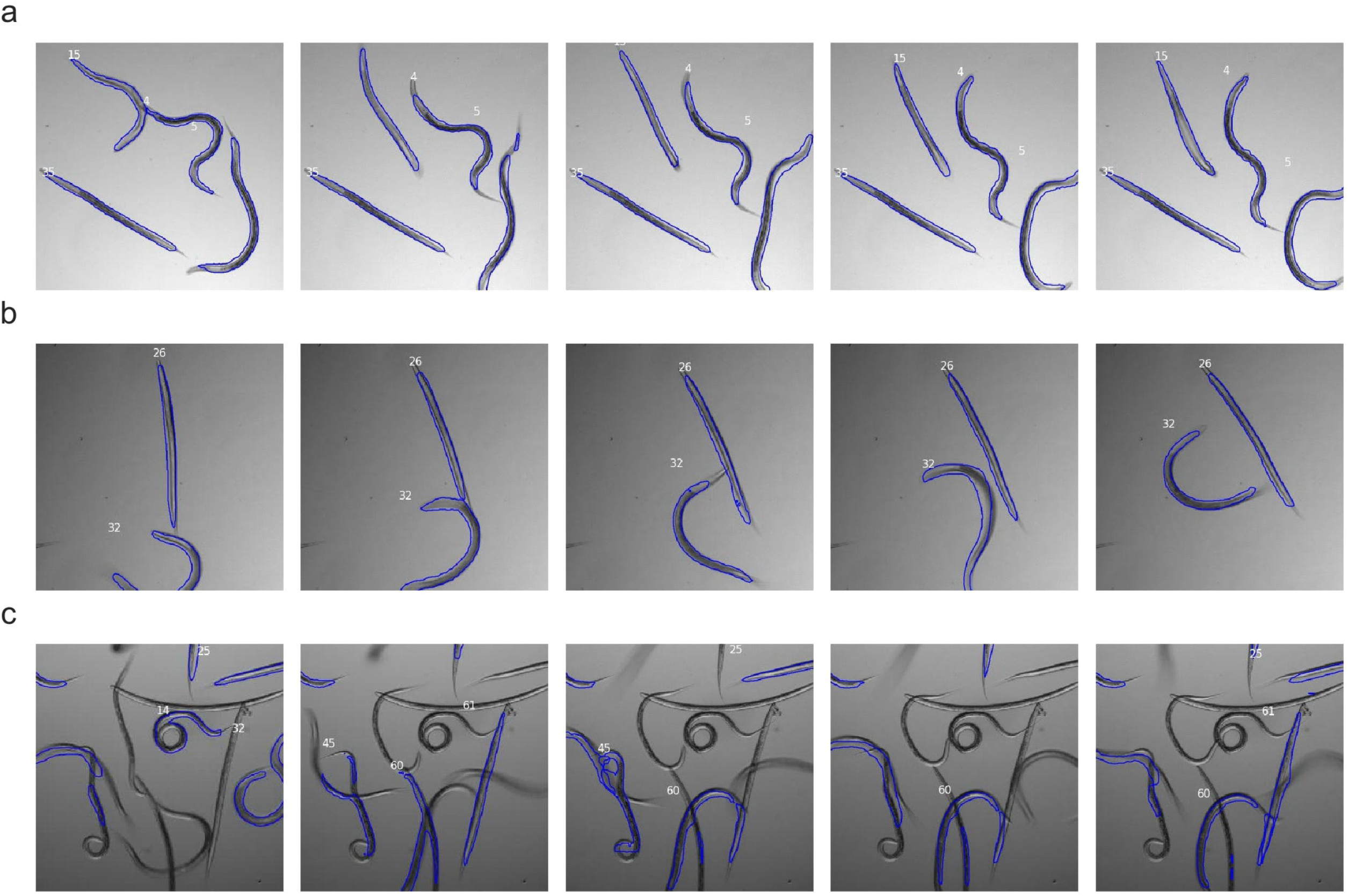
Sample frames of common detection scenarios in motility videos. The blue outlines are the masks detected by Mask R-CNN. Each detected worm has an associated identification number displayed in white based on the tracking across the frames. **(a) c**ommon scenario with low or no overlaps of individual worms. **(b) e**xample of a motile worm (id 32) colliding with a non-motile worm (id 26) causing the later to be perceived as motile. **(c) c**omplex detection scenario with a high number of worms with significant overlaps

### 3.2. Motility forecast errors

In the manual video processing, we observed the motility and the live rate. To measure the performance of the algorithms, we used the motility rate rather than the live rate even though the stillness alone does not denote a dead worm. The reason for this choice was that the algorithms detect movement and are not able to differentiate if the movement was caused by the object itself or by an interaction with another object. The general manual motility rate was always higher than the manual live rate (Supplementary Table S1). We measured the performance of the individual algorithms by evaluating the errors, defined as the differences between the manual processing of the file and the respective algorithm. Out of the three methods, Mask R-CNN had the lowest MAE with a value of 5.6%, followed by WF-NTP (8.76%) and the WI (14.2%). In terms of the bias of the predictions, the WI had the lowest ME of -0.71%, therefore slightly overestimating the motility rate on average, while both WF-NTP and Mask R-CNN underestimated the motility rate with values of 6.52% and 1.95% respectively (Table 1).

**Table 1:** Motility error metrics for algorithms

| Algorithm | MAE, % | ME, % |
| --- | --- | --- |
| Wiggle Index | 14.2 | -0.71 |
| WF-NTP | 8.76 | 6.52 |
| Mask R-CNN | 5.61 | 1.95 |

Shown in Fig 3, the WI had an increasing volatility with an increase in the ratio of alive worms in the group. Mask R-CNN overestimated the motility rate for the motility group 0; while four videos had a zero motility rate, showing that the overestimation is not systematic, two videos had a motility rate error below 3.6%, and the remaining three videos had high motility rates without any apparent reason. WF-NTP and Mask R-CNN had a tendency to systematically underestimate for the motility group 100 (Fig 3). The mean and standard deviation of motility rates per motility group and algorithm is shown also in Supplementary Table S2 and the same statistics are reported as differences between the manual count and the respective algorithm in Supplementary Table S3.

**Fig 3:**
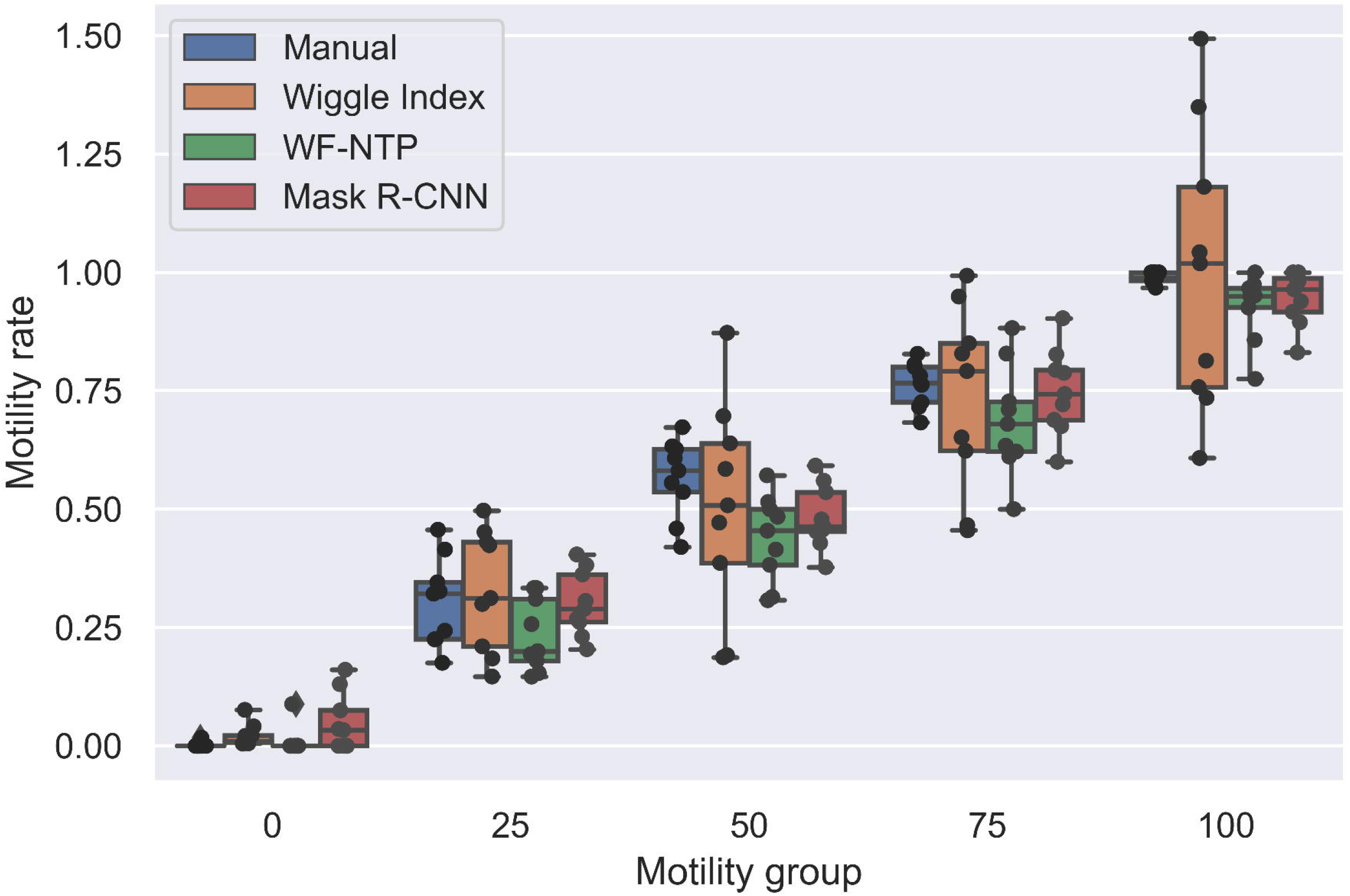
Motility rates comparisons for individual motility groups per algorithm. The number of the motility group indicates the percentage of live larvae. The motility rate is visualized as box plots.

The differences between the manual processing and the respective algorithm were expressed as the mean absolute error (MAE) and mean error (ME).

### 3.3. Instance detection performance

The motility rates are an indicator of the degree of correctly classified worms as motile or non-motile based on IoU. However, they do not provide the information of the number of detected worm instances in the image. In order to ensure that the motility rates were estimated reliably and not only based on a small subset of detected worms, the detection performance was validated. To achieve this, we compared Mask R-CNN and WF-NTP for the total worm count (WI was excluded as the algorithm does not detect individual instances of worms, therefore this information was not available). The precision was measured based upon the error term between the manual counts and the respective algorithm. In terms of the size of the error, the MAPE was used and we observed that the error was significantly lower (P < 0.0001, Wilcoxon signed-rank test) for Mask R-CNN (7.6%) than for WF-NTP (40.23%). Both algorithms had a tendency to underestimate the number of worms on average, therefore not detecting all the worm instances in a video, mainly due to the challenges of detecting overlapping or clustered worms. This was confirmed by the MPE with values of 5.61% for Mask R-CNN and 40.23% for WF-NTP (Table 2). The reported MAPE and MPE for WF-NTP had the same value because all the detections underestimated the total number of worms. WF-NTP underestimated the worm count for all motility groups, while Mask R-CNN had similar box plots in comparison to the manual counts apart from underestimating the motility group 0, as shown in Fig 4. The mean error and standard deviation of the differences between the manual count and the respective motility algorithm is shown in Supplementary Table S4.

**Table 2:** Worm count error metrics for algorithms

| Algorithm / Metric | MAPE, % | MPE, % |
| --- | --- | --- |
| WF-NTP | 40.23 | 40.23 |
| Mask R-CNN | 7.6 | 5.61 |

**Fig 4:**
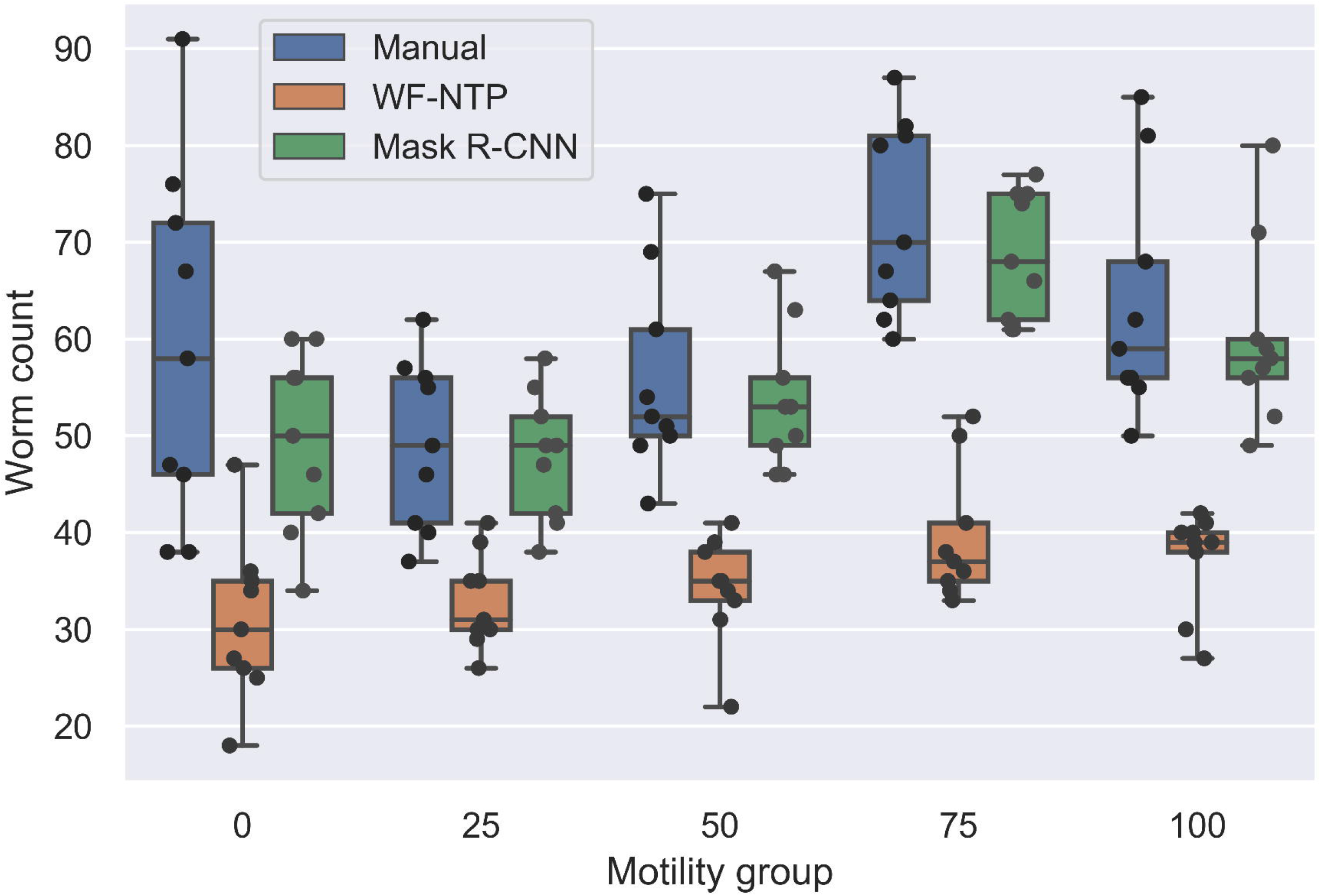
Worm count comparisons for individual motility groups per algorithm. The number of the motility group indicates the percentage of live larvae. The number of worms is visualized in box plots.

The differences between the manual counts and the respective algorithm were expressed as the mean absolute percentage error (MAPE) and mean percentage error (MPE).

### 3.4. Mask R-CNN detection and classification precision

The precision of detecting worm instances in a video was on average 98.6%. This was calculated based on all the tracked worms across all the motility groups without setting a minimum number of frames for which a worm needed to be tracked. The detections were manually validated and the detection was considered correct despite only a part of the worm being detected in some frames. The majority of cases where the detection was incorrect was due to debris that had a similar size and shape as a worm. For the detected worms across all the motility groups, there was an overall accuracy of 89% to correctly label them as either motile or non-motile based on their mean IoU values. In Fig 5, the distribution for different IoU values obtained from all the worms across all the motility groups clearly showed that most of misclassified cases were located around the 0.8 threshold, while the classification accuracy increased for values further away from the threshold. The precision and recall values (Table 3) confirm that the prevalent case was the misclassification of motile worms as non-motile.

**Table 3:** Classification performance metrics

| Classification | Precision, % | Recall, % |
| --- | --- | --- |
| Motile | 94 | 88 |
| Non-Motile | 84 | 92 |

**Fig 5:**
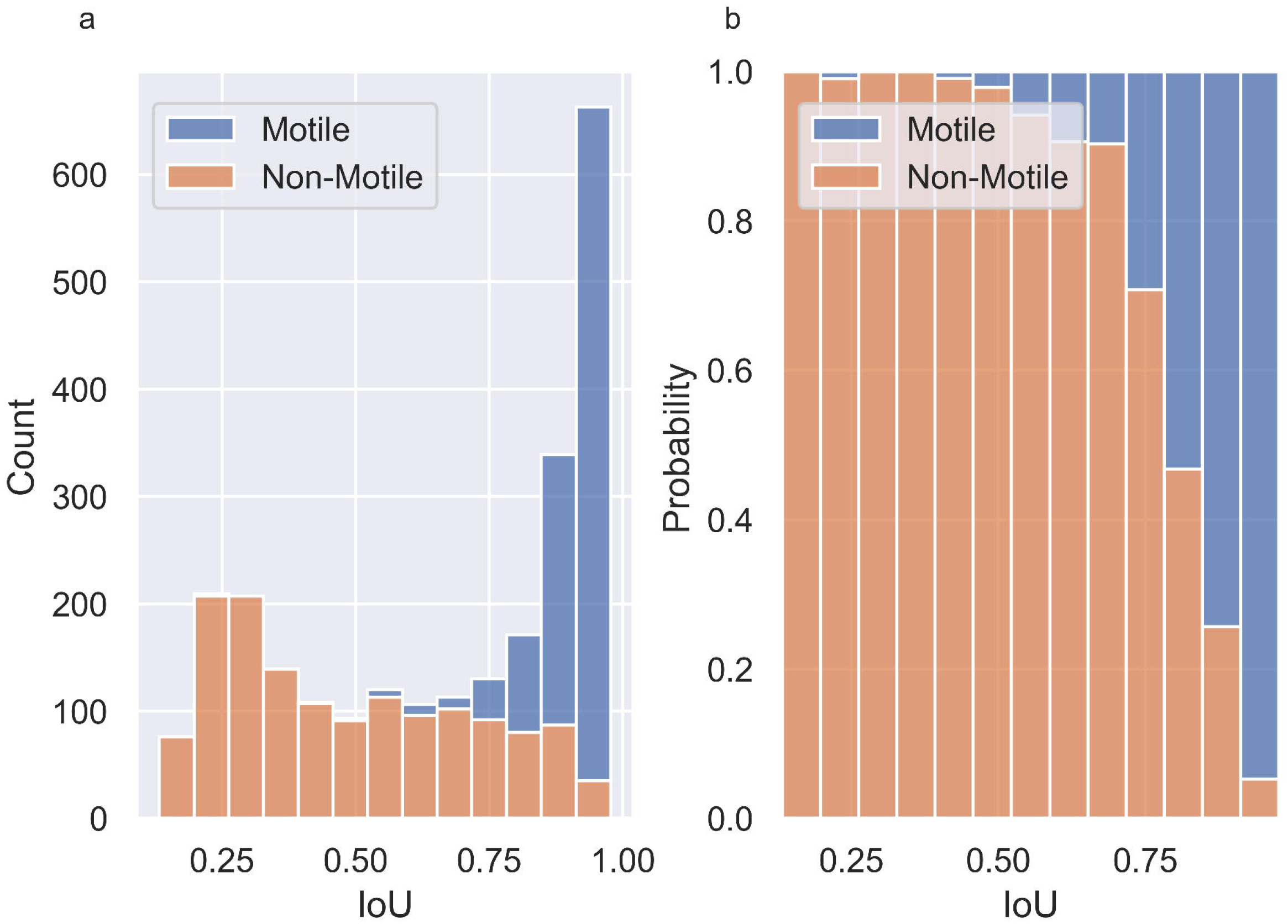
The distribution for different Intersection over Union (IoU). Both **(a) a** histogram of IoU and the associated number of worms per bin classified into motile / non-motile and (**b)** a histogram of IoU and the associated probability per class (motile / non-motile) show that the majority of misclassified cases were located around the 0.8 IoU.

The errors defined as differences between Mask R-CNN and the manually evaluated motility rates were tested for normality using the D’Agostino-Pearson test as well as Shapiro-Wilk test. In both cases we could not reject the null hypothesis that the data is from a normal distribution with p-values 0.85 and 0.35 respectively (Table 4).

**Table 4:**
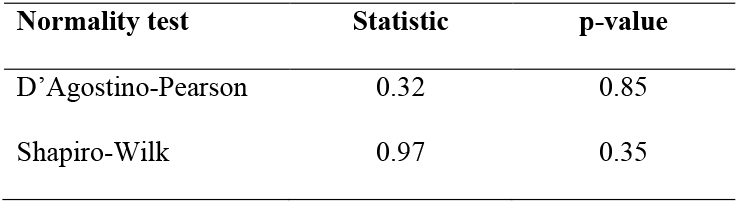
Normality tests for the motility rate error terms between Mask R-CNN and the manual processing

| Normality test | Statistic | p-value |
| --- | --- | --- |
| D'Agostino-Pearson | 0.32 | 0.85 |
| Shapiro-Wilk | 0.97 | 0.35 |

Precision and recall metrics for classification performance

For both normality tests we cannot reject the null hypothesis that the data is from a normal distribution.

### 3.5. Monte Carlo drug screening simulation

The Monte Carlo simulation was based on the results from the motility forecasts and the obtained forecast errors. While WF-NTP and Mask R-CNN had a positive correlation of forecast errors (0.65), the forecast errors for WI were not correlated with the other algorithms (Table 5).

**Table 5:** Correlation matrix for the prediction errors of the individual algorithms

|  | Wiggle Index | WF-NTP | Mask R-CNN |
| --- | --- | --- | --- |
| Wiggle Index | 1 |  |  |
| WF-NTP | -0.04 | 1 |  |
| Mask R-CNN | 0.08 | 0.65 | 1 |

Mask R-CNN achieved the highest accuracy in the simulation with a value of 94% followed by WF-NTP 90% and the WI 85% (full classification performance in Supplementary Table S5). The simulation estimated that an improved detection performance of 3.15% (MAE) between Mask R-CNN and WF-NTP led to a 3.5% improvement of drug screening accuracy, while an improvement of 8.59% (difference of MAE between Mask R-CNN and WI) led to a 9% improvement of drug screening accuracy.

## 4. Discussion

The motility phenotype is often used for drug screening and drug testing of anthelmintics. In our study, we deployed and compared three algorithms to measure motility in the parasitic nematode *H. contortus*. Apart from evaluating motility for both WF-NTP and Mask R-CNN, we were also interested in the detection precision of worms by these algorithms. The reason for that was the fact that for both algorithms, the detection was the first step followed by the tracking across the frames and the classification into motile and non-motile. Therefore, accurate detection was the pre-requisite for reliable motility quantification.

The results showed that the performance increased with an increased complexity of algorithm. The WI, despite requiring minimal configuration and being the fastest algorithm, had some limitations in comparison to the other methods; it was by design not able to detect individual instances of worms and a control group with only motile worms needed to be created in order to be able to translate the WI into a percentage for the motility rate for each video, otherwise it was hard to interpret the numerical value of the WI. Because of these limitations and the lack of precision, it would be advisable to use WI only for initial studies to quickly test experimental results.

For WF-NTP, overlapping parasites had the highest negative impact on the detection ability, as the algorithm filters out objects that are above or below a certain pixel size. This resulted in detecting overlapping worms as a single object and these objects were then excluded based on the size threshold. As a consequence, the total number of detected instances in an image was significantly lower than the actual number obtained by manually processing the videos. Although WF-NTP struggled with detecting all instances, the classification of motile / non-motile performed well once an instance was detected.

Mask R-CNN had the lowest MAE. However, similarly to WF-NTP, it struggled with overlapping objects, primarily due to the fact that the algorithm uses NMS. As a result, some instances were not detected, although in contrast to WF-NTP, the worm clusters were not discarded as a whole. The problem of detecting overlapping objects is currently an area of active research in object detection, particularly for biological data. For instance, Böhm et al. [35] are working on algorithms that can handle overlapping objects and have had success with applying their algorithm to *C. elegans*. Motile worms had a higher chance of being detected, because they had higher variability across the frames due to their movement. In terms of precision, the algorithm was slightly biased towards underestimating the motility rate; however, the errors were normally distributed enabling us to increase the number of acquired videos in order to get to a desired level of confidence and therefore successfully replace manual processing.

The results confirmed that IoU could be used as a metric for determining the motility of larvae, although some modifications might be required in order to deal with slow-moving worms, such as including the total distance travelled during the video to complement the frame-by-frame information. Furthermore, the IoU threshold should be considered a parameter to be optimized as it can vary based on the age of the larvae, the temperature of the environment and other factors. The downside of using Mask R-CNN is that it requires a quite big amount of annotated data (provided in the Data availability), which is time consuming to create and the training runs for 1-2 days on a GPU instance (an Amazon Web Services p2.xlarge compute instance with 1 GPU, 4 vCPUs and 61GB RAM was used).

The drug screening simulation showed that the improved detection performance resulted in increased accuracy for drug screening of roughly the same magnitude as the MAE improvement, illustrating that the improvement is not negligible for practical purposes even though some of the forecast errors of the algorithms were correlated. For applications with multiple observations for a given category / group, the impact would be most likely lower, because with an increase in the number of observations per category / group, the sample mean of the forecast errors would converge to the expected value of the forecast errors. In this situation, the bias of the estimate measured in our case by ME would then have a higher impact than the variance which would make WF-NTP, with the highest bias, the least accurate algorithm.

In general, for quick low-precision experiments, such as primary screening of large compound libraries, the best option is to use the WI as it can be used out-of-the-box with slight modifications to a few parameters. If the detection of individual instances is desired, Mask R-CNN is the best choice, although it is more time-intensive to prepare the data and train the model. Moreover, the ability to precisely detect larvae in an image then allows the researchers to perform multiple types of analyses that involve microscopy of the parasites and are not only limited to motility assays, but also include other common *in vitro* assays for the diagnosis of drug resistance, e.g. larval development tests and egg hatch tests [36]. In the aspect of potentially repurposing the algorithms for other assays, Mask R-CNN is the most flexible of the models, because it is able to detect multiple classes of objects. On contrary, WF-NTP decides solely on the size of the objects, making a distinction between classes of similar sizes impossible.

## 5. Conclusions

Our study confirmed that the state-of-the-art machine learning algorithm Mask R-CNN was able to outperform less computationally complex algorithms that are based on pixel intensity or Gaussian threshold. The algorithm performed better in the detection of individual worm instances as well as the subsequent motility classification. This was consistent with how the algorithm is designed as the tracking and motility classification depend on the detection accuracy. For the motility classification, we used IoU as a new metric, that successfully replaced previous ones based on locomotion specific to the studied organisms. Whilst IoU is very flexible, as it only requires a single parameter, it would need further extension to be able to deal with cases of heterogeneous groups of viable worms that have a high variance in their speed of movement.

The gain in precision of Mask R-CNN came at the cost of requiring an extensive annotated dataset as well as computational resources for training the model. Therefore, the time-precision trade-off needs to be evaluated by the researchers on a case-by-case basis, unless the study allows reapplying an already existing trained model. The output of the present study provides an annotated dataset containing various developmental stages of *H. contortus* that can be utilised by researchers for future machine learning applications.

## Data availability

All underlying data for training the model, the trained model itself as well as the input and output of the motility analysis for all three algorithms is available at https://doi.org/10.5281/zenodo.5734143. The source code for Mask R-CNN is available in a repository on GitHub https://github.com/zofkam/mask_rcnn_motility. We have also used Zenodo to assign a DOI to the repository https://doi.org/10.5281/zenodo.5735068.

## Additional files

**Supplementary Table S1**: Manual counts of motility and live rates per motility group

**Supplementary Table S2**: Motility rates per motility group and algorithm

**Supplementary Table S3:** Mean error between the manual count and the respective motility algorithm

**Supplementary Table S4**: Mean of differences in worm counts between the manual processing and the respective algorithm

**Supplementary Table S5:** Drug screening simulation classification performance

**Supplementary Fig S1**: Average number of worms in the videos per motility group for WF-NTP and Mask R-CNN.

**Supplementary Fig S2**: The impact of the length of the video in frames on the percentage error of the detected worms for Mask R-CNN.

**Supplementary Video**: Mask R-CNN sample detection video of motility group 50.

### Abbreviations

CNN: convolutional neural network
fps: frames per second
GPU: graphics processing unit
IoU: intersection over union
L3: third-stage larva
MAE: mean absolute error
mAP: mean average precision
MAPE: mean absolute percentage error
Mask R-CNN: region based convolutional neural network
ME: mean error
MPE: mean percentage error
NMS: non-maximum suppression
ROI: region of interest
RPN: regional proposal network
WF-NTP: Wide Field-of-View Nematode Tracking Platform
WI: Wiggle Index

## Competing Interests

The authors declare that they have no competing interests.

## Funding

This work was supported by Charles University UNCE/18/SCI/012 and SVV 260 550 as well as by the project EFSA-CDN [CZ.02.1.01/0.0/0.0/16_019/0000841], co-funded by ERDF.

## Authors’ contributions

M.Ž. and L.T.N. designed this study and wrote the manuscript. M.Ž. implemented the deep learning model, wrote the code, ran the statistical analyses and generated the figures. L.T.N. created the graphical abstract. L.T.N., E.M. acquired and annotated the images used to train the deep learning model. M.Ž. and L.T.N. conducted the experiments and analyzed the results. P.M. managed the project. All authors contributed to literature review and critical revision of the manuscript.

## Acknowledgements

We thank MVDr. Tomáš Kaňka for assisting with the sheep infection. The authors thank prof. Lenka Skálová for critical discussion and comments. We would like to express our gratitude to Jon Nolan for proofreading and revisions.

## Notes

### Competing Interest Statement

The authors have declared no competing interest.

### Summary of Updates

The manuscript has been thoroughly revised, the Figures renumbered and two figures added. Supplemental video added.

