## Supplementary material for "Image recognition based on deep learning in *Haemonchus contortus* motility assays": Supplemntary files: Supplementary_files - revisions.docx

**Supplementary Table S1**: Manual counts of motility and live rates per motility group

| Motility group | Live rate | Motility rate |
| --- | --- | --- |
| 0 | 0.2 ± 0.6 | 0.2 ± 0.6 |
| 25 | 26.3 ± 7.3 | 30.4 ± 9.4 |
| 50 | 50.3 ± 8.7 | 56.6 ± 8.3 |
| 75 | 72.3 ± 6.4 | 76.3 ± 4.7 |
| 100 | 98.2 ± 1.2 | 99 ± 1.1 |

The data of manual counts are expressed as mean (%) ± SD (%). Motility rate was higher than the live rate because the collisions of live worms propelled dead worms causing them to be motile.

**Supplementary Table S2**: Motility rates per motility group and algorithm

| Motility group | Manual | Wiggle Index | WF-NTP | Mask R-CNN |
| --- | --- | --- | --- | --- |
| 0 | 0.2 ± 0.6 | 2.2 ± 2.3 | 1.0 ± 2.9 | 4.8 ± 6.1 |
| 25 | 30.4 ± 9.4 | 32.8 ± 12.8 | 23.4 ± 7.6 | 30.1 ± 6.9 |
| 50 | 56.6 ± 8.3 | 50.4 ± 22.7 | 43.8 ± 9.1 | 48.3 ± 6.8 |
| 75 | 76.3 ± 4.7 | 73.4 ± 19.6 | 68.8 ± 11.6 | 74.9 ± 9.0 |
| 100 | 99.0 ± 1.1 | 100.0 ± 30.0 | 92.8 ± 7.0 | 94.6 ± 5.7 |

The data are expressed as mean (%) ± SD (%).

**Supplementary Table S3**: Mean error of the differences between the manual count and the respective motility algorithm

| Motility group | Wiggle Index | WF-NTP | Mask R-CNN |
| --- | --- | --- | --- |
| 0 | 2.0 ± 1.9 | 0.8 ± 2.4 | 4.6 ± 5.7 |
| 25 | 2.5 ± 12.3 | 0.0 ± 12.3 | 0.0 ± 8.4 |
| 50 | -6.2 ± 21.7 | -12.8 ± 11.4 | -8.3 ± 5.5 |
| 75 | -2.9 ± 19.8 | -7.5 ± 9.5 | -1.4 ± 4.8 |
| 100 | 1.0 ± 29.5 | -6.2 ± 7.0 | 0.0 ± 5.3 |

The data are expressed as mean error (%) ± SD (%).

**Supplementary Table S4**: Mean of differences in worm counts between the manual processing and the respective algorithm

| Motility group | WF-NTP | Mask R-CNN |
| --- | --- | --- |
| 0 | -28.3 ± 14.4 | -9.9 ± 10.8 |
| 25 | -16.3 ± 9.2 | -1.3 ± 3.3 |
| 50 | -21.8 ± 8.3 | -2.3 ± 4.6 |
| 75 | -33.0 ± 8.2 | -3.8 ± 4.2 |
| 100 | -26.2 ± 11.1 | -3.3 ± 4.4 |

The number worms in each video was counted manually and also accessed by algorithm. The differences were averaged, and the data are expressed as mean ± SD.

**Supplementary Table S5**: Drug screening simulation classification performance

|  | Wiggle Index | WF-NTP | Mask R-CNN |
| --- | --- | --- | --- |
| Weighted Precision | 85 | 90 | 94 |
| Weighted Recall | 85 | 91 | 94 |
| Accuracy | 85 | 90 | 94 |

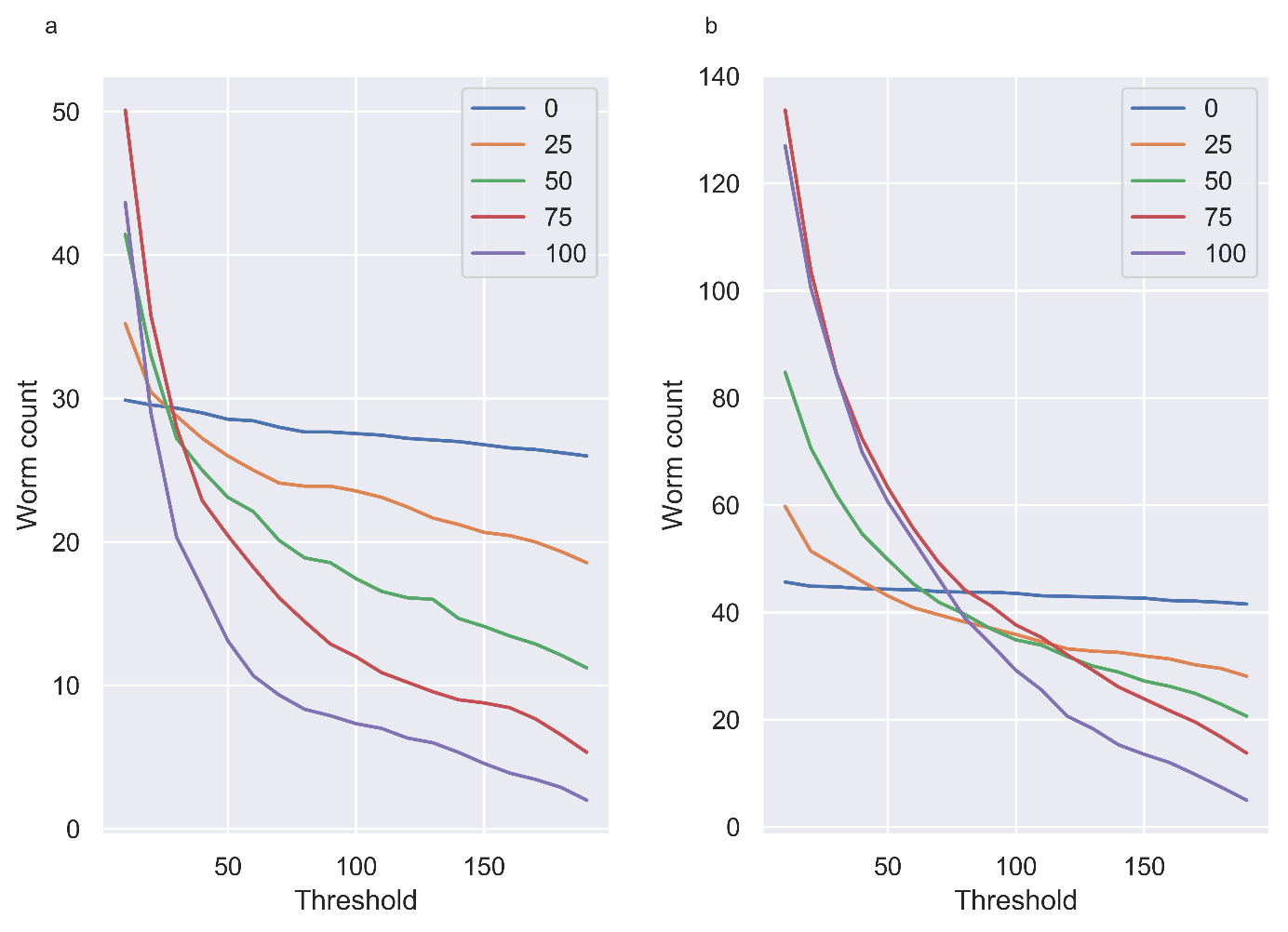

**Supplementary Figure S1**: Average number of worms in the videos per motility group for WF-NTP and Mask R-CNN. The graph for **(a)** WF-NTP and **(b)** Mask R-CNN shows how the number of detections changes as we increase the minimum number of frames that a worm needed to be detected for to be counted. The higher the motility group the more duplicate detections occurred, this was caused by a higher rate of overlaps and movement in motile groups. The plot showed the need to limit the number of detections to the maximum number of instances detected in a single frame over the duration of the video.

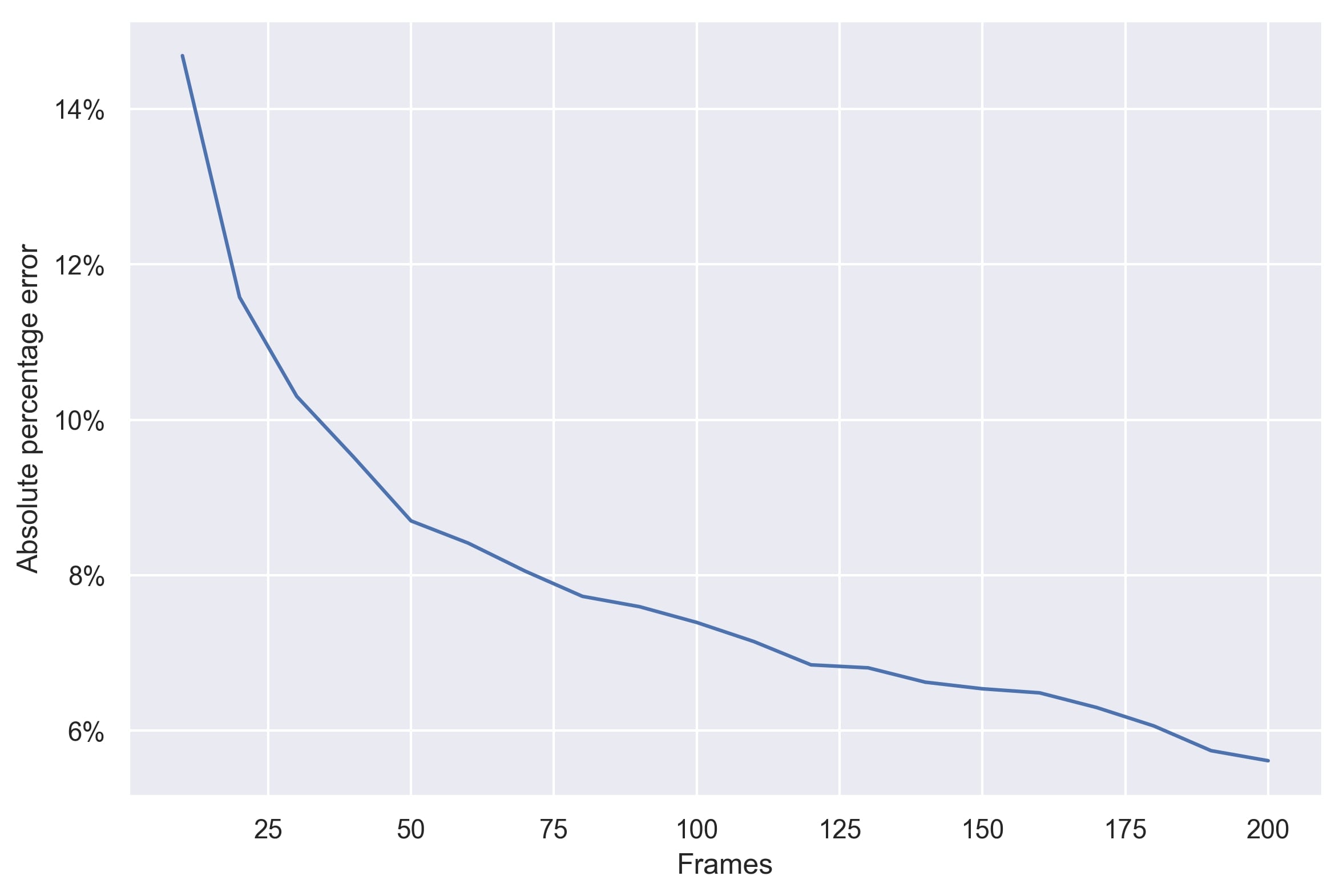
**Supplementary Figure S2**: The impact of the length of the video in frames on the absolute percentage error of the detected worms for Mask R-CNN. With an increasing length of the videos the absolute error decreases.
