## Supplementary figures and images for "Image recognition based on deep learning in *Haemonchus contortus* motility assays"

### Supplementary_fig1.jpg

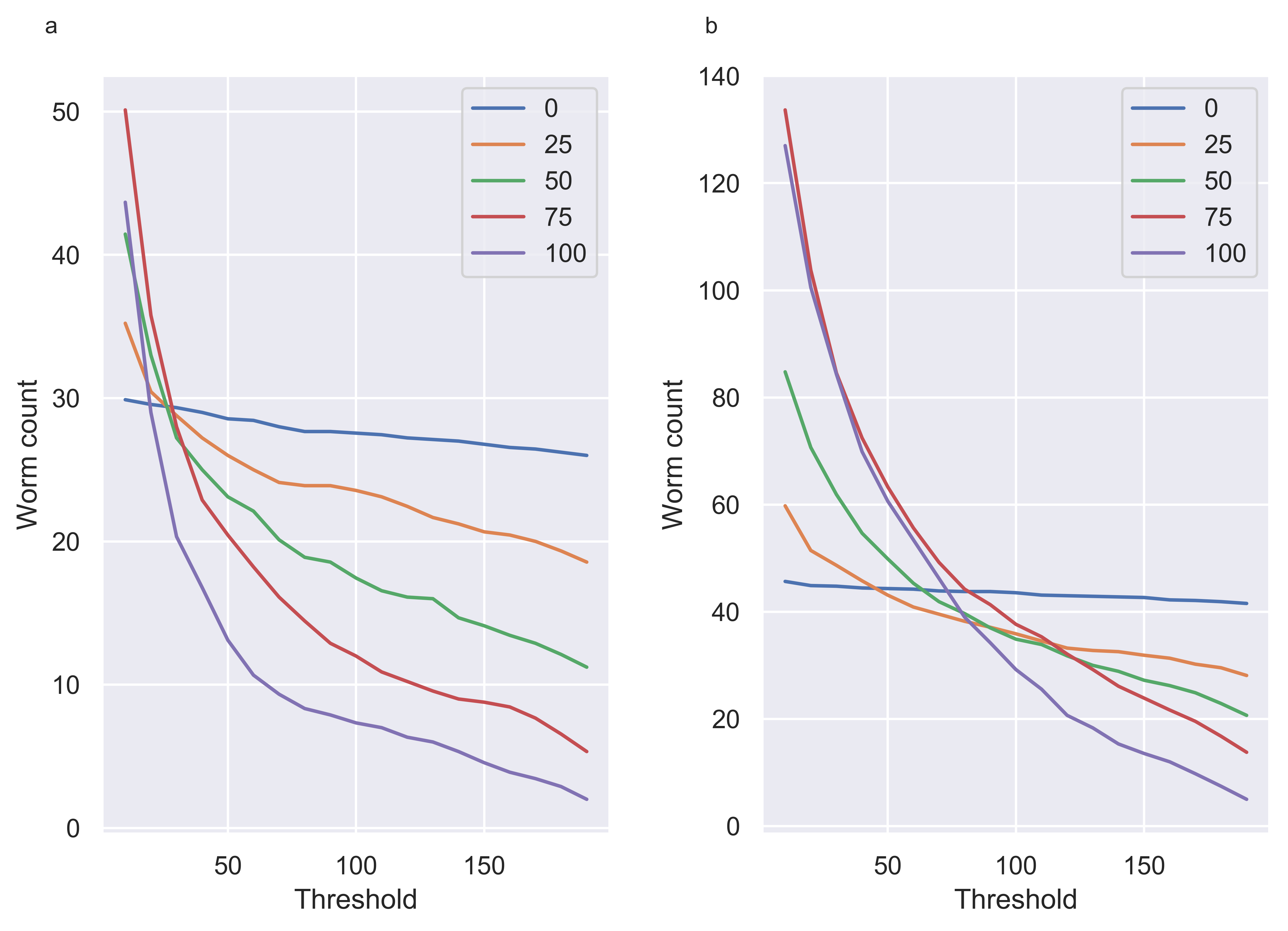

### Supplementary_fig2.jpg

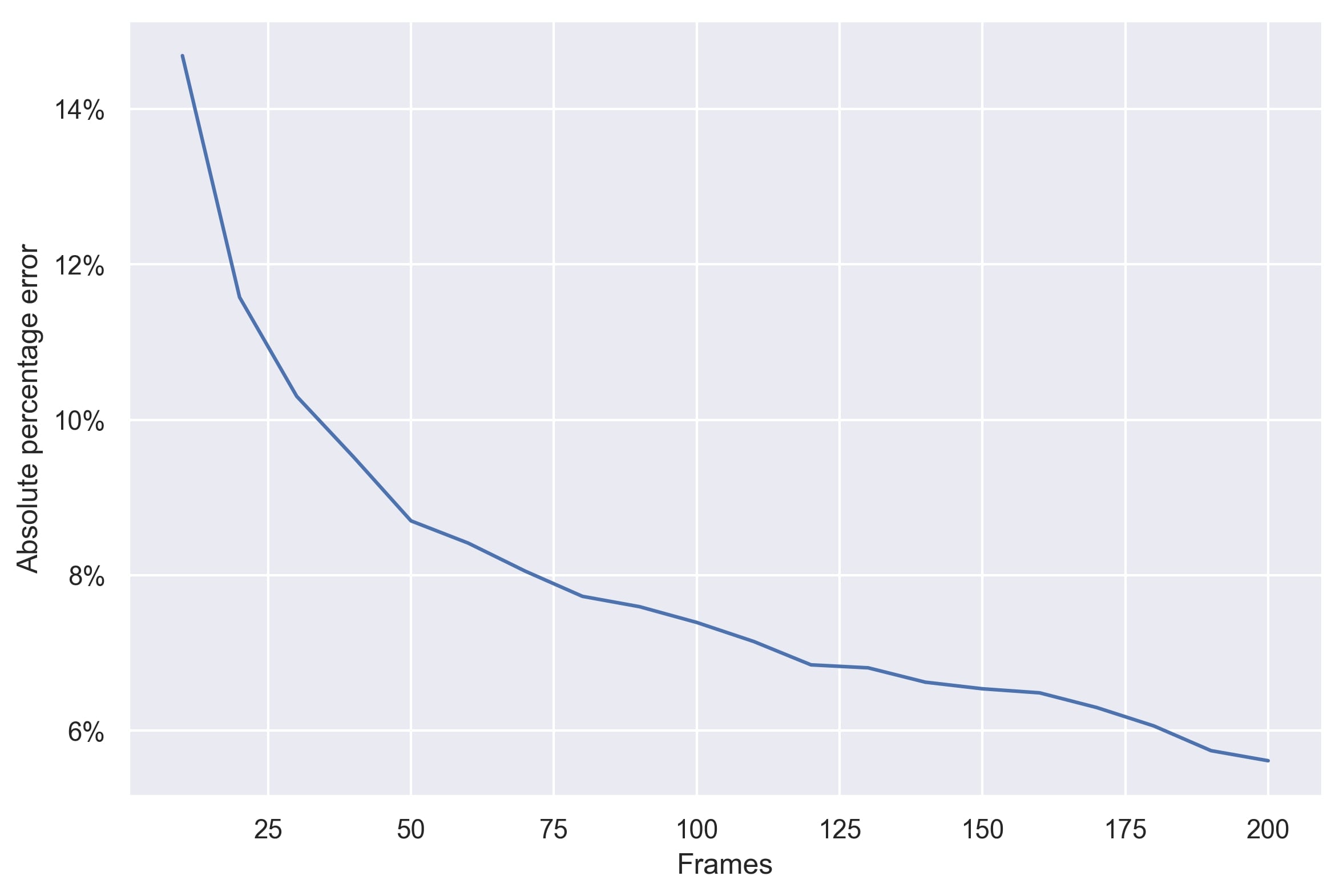
